# Stomatal patterning is differently regulated in adaxial and abaxial epidermis in Arabidopsis

**DOI:** 10.1101/2024.02.22.581564

**Authors:** Pirko Jalakas, Ingmar Tulva, Nele Malvīne Bērziņa, Hanna Hõrak

## Abstract

Stomatal pores in leaves mediate CO_2_ uptake into the plant and water loss via transpiration. Most plants are hypostomatous with stomata present only in the lower leaf surface (abaxial epidermis). Many herbs, including the model plant *Arabidopsis thaliana*, have substantial numbers of stomata also on the upper (adaxial) leaf surface. Studies of stomatal development have mostly focused on abaxial stomata and very little is known of adaxial stomatal formation. We addressed the role of leaf number in determination of stomatal density and stomatal ratio, and studied adaxial and abaxial stomatal patterns in mutants deficient in known abaxial stomatal development regulators. We found that stomatal density in some genetic backgrounds varies between different fully expanded leaves and recommend using defined leaves for analyses of stomatal patterning. Our results indicate that stomatal development is at least partly independently regulated in adaxial and abaxial epidermis, as i) plants deficient in ABA biosynthesis and perception have increased stomatal ratios, ii) the *epf1epf2*, *tmm* and *sdd1* mutants have reduced stomatal ratios, iii) *erl2* mutants have increased adaxial but not abaxial stomatal index, and iv) stomatal precursors preferentially occur in abaxial epidermis. Further studies of adaxial stomata can reveal new insights into stomatal form and function.

## Introduction

Stomatal pores in plant leaves mediate CO_2_ uptake for photosynthesis and water loss via transpiration. Stomatal numbers and size determine the maximal potential for stomatal conductance, a trait often positively related with yield (Roche, 2015). At the same time, lower stomatal densities are associated with increased water use efficiency and drought tolerance (Hughes *et al*., 2017; Caine *et al*., 2019; Dunn *et al*., 2019). Hence, understanding how stomatal numbers are determined in leaves can help in breeding for varieties with increased productivity or water use efficiency.

Knowledge on stomatal formation in dicot leaves mostly originates from studies of abaxial epidermal development in the model plant *Arabidopsis thaliana* (Arabidopsis); key stomatal development pathway components are shown in Figure 1. Stomatal differentiation is achieved via sequential activation of the transcription factors SPEECHLESS (SPCH, MacAlister *et al*., 2007; Pillitteri *et al*., 2007), MUTE (MacAlister *et al*., 2007; Pillitteri *et al*., 2007) and FAMA (Ohashi-Ito and Bergmann, 2006). The central differentiation path is regulated by cell-to-cell signaling via peptides (epidermal patterning factors, EPFs) perceived by cell surface receptors to ensure optimal stomatal spacing. EPF1 and EPF2 peptides (Hara *et al*., 2007; Hunt and Gray, 2009) are perceived by a membrane complex formed by a leucine-rich repeat receptor-like kinase from the ERECTA (ER) family (Shpak *et al*., 2005) and the receptor-like protein TOO MANY MOUTHS (TMM) (Yang and Sack, 1995; Lee *et al*., 2012; Lin *et al*., 2017). The ER family includes the ER, ERECTA-LIKE 1 (ERL1), and ERL2 proteins that have partly overlapping and partly distinct functions (Shpak *et al*., 2005). ER is considered a major receptor for EPF2 and ERL1 for EPF1 (Lee *et al*., 2012). The binding of EPF1 or EPF2 to the receptor complex activates downstream signaling that ultimately suppresses stomatal differentiation. Cleavage of pro-peptides is necessary to form biologically active signaling peptides; an apoplastic subtilisin-like serine protease STOMATAL DENSITY AND DISTRIBUTION 1 (SDD1) that suppresses stomatal development has been suggested to carry out this function (Berger and Altmann, 2000; Von Groll *et al*., 2002). Loss-of-function mutations in *EPF1* and *EPF2*, *ER*-family genes, *TMM,* or *SDD1* lead to increased stomatal densities in Arabidopsis.

**Figure 1.**
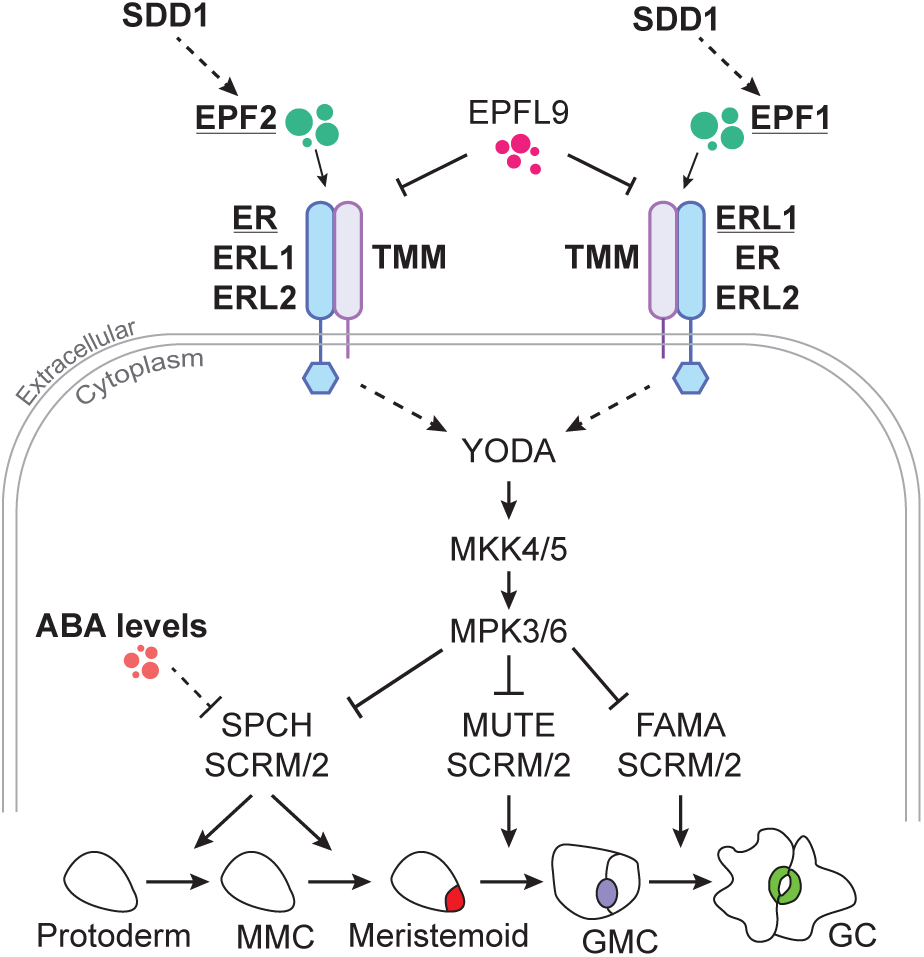
Schematic model of stomatal development pathway in Arabidopsis. The subtilisin-like serine protease SDD1 is suggested to process the cleavage of pro-peptides to form biologically active signaling peptides, e.g. EPF1 and EPF2. EPF1/2 (and the antagonistic EPFL9 (STOMAGEN) that competes with EPFs) bind to the cell-surface receptor complex of a member of the ERECTA family (ER, ERL1, or ERL2) and TMM. ER is considered a major receptor for EPF2, and ERL1 for EPF1. Binding to the membrane complex activates the downstream signaling mitogen-activated protein kinase (MAPK) cascade composed of YODA–MKK4/5–MPK3/6 by a yet unidentified mechanism. The activated MAPK cascade represses stomatal production through three bHLH transcription factors (SPCH, MUTE, and FAMA) and their partner bHLH proteins SCREAM (SCRM) and SCRM2. ABA is thought to suppress stomatal production through SPCH. Key regulators studied in this work are shown in bold. Meristemoid mother cell (MMC); guard mother cell (GMC); guard cell (GC).

Abscisic acid (ABA) is a phytohormone produced in response to dry air or soil water deficit in plants (McAdam *et al*., 2016; Kuromori *et al*., 2022). Recently, ABA was shown to suppress stomatal development in Arabidopsis via the SUCROSE NONFERMENTING 1-related protein kinases SnRK2.2, SnRK2.3, and SnRK2.6 (OST1) that phosphorylate SPCH, leading to its degradation (Yang *et al*., 2022a). ABA biosynthesis-deficient mutants *nced3nced5*, *aba2-2*, and *aba3-1* have higher stomatal densities or indices (Tanaka *et al*., 2013*a*; Chater *et al*., 2015; Jalakas *et al*., 2018), while the ABA catabolism mutant *cyp707a1cyp707a3* has lower stomatal numbers (Tanaka *et al*., 2013*a*; Jalakas *et al*., 2018).

While most plants are hypostomatous with stomata located only in the abaxial epidermis (Salisbury and Oliver, 1928; Muir, 2015), Arabidopsis is an amphistomatous plant that produces a substantial proportion of its stomata in the adaxial epidermis (Berger and Altmann, 2000; Pantin *et al*., 2013; Hronková *et al*., 2015; Vráblová *et al*., 2017; Tulva *et al*., 2023, Preprint; Watts *et al*., 2024). Relatively little is known of the mechanisms of adaxial stomatal development and whether it is regulated by the same components and in a similar manner as stomatal formation in the abaxial epidermis. Previous studies suggest that adaxial and abaxial stomatal development are at least partly independently regulated. For example, stomatal density and index in the adaxial epidermis seem to be more responsive to changes in light conditions and relative air humidity (Hronková *et al*., 2015; Devi and Reddy, 2018; Tulva *et al*., 2023, Preprint). Stomatal ratio is altered in some Arabidopsis mutants lacking or overexpressing known regulators of stomatal development (Berger and Altmann, 2000; Kong *et al*., 2012; Dow *et al*., 2014; Franks *et al*., 2015; Hronková *et al*., 2015; Vráblová *et al*., 2017; Qi *et al*., 2019), pointing at differences between stomatal developmental mechanisms in the upper and lower leaf surface.

Here we address the within-plant variation of stomatal density and ratio in Arabidopsis and use mutants deficient in known regulators of stomatal development to test whether these signaling components similarly affect adaxial and abaxial stomatal development. We also analyze mutants with different ABA levels to understand if ABA affects adaxial and abaxial stomatal formation in a similar manner.

## Materials and Methods

### Plant lines used in experiments

*Arabidopsis thaliana* accession Col-0 (wild type) and mutants in the same background were used for experiments, details for mutants used in experiments are listed in Table 1. T-DNA insertion mutants were from the SALK (Alonso *et al*., 2003) and GABI-Kat collections (Rosso *et al*., 2003) and ordered from Nottingham Arabidopsis Stock Centre (NASC, (Scholl *et al*., 2000)). Primers used for genotyping studied mutants are shown in Supplementary Table 1. We note that the *tmm* T-DNA insertion line SALK_115723C used in our study also harbors a T-DNA insertion in *STKR1* (Nietzsche *et al*., 2018). Although this line is a *tmm stkr1* double mutant, its stomatal phenotypes are similar to previously published results for other *tmm* alleles, hence we refer to the mutant as *tmm*.

**Table 1.** Mutants used in experiments.

| **Mutant name** | **Mutant short name** | **Mutation/T-DNA insertion line name** | **Reference** | **Expected phenotype** | **Observed adaxial SD** | **Observed abaxial SD** | **Observed stomatal ratio (SR)** |
| --- | --- | --- | --- | --- | --- | --- | --- |
| *nced3-2nced5-2* | *nced3/5* | GK-129B08, GK-328D05 | (Frey *et al.*, 2012) | low ABA levels, increased SD | increased SD | increased SD | increased SR |
| *epf1-1epf2-2* | *epf1/2* | SALK_137549, GK-673E01 | (Hunt and Gray, 2009) | increased SD | increased SD | increased SD | decreased SR |
| *nced3-2nced5-2 epf1-1epf2-2* | *nced3/5 epf1/2* | GK-129B08, GK-328D05, SALK_137549, GK-673E01 | created by crossing *nced3-2nced5-2* and *epf1-1epf2-2* in this study | low ABA levels, increased SD | increased SD | increased SD | normal SR |
| *cyp707a1-1 cyp707a3* | *cyp707a1/a3* | SALK_069127, SALK_101566 | (Kushiro *et al.*, 2004; Okamoto *et al.*, 2006) | high ABA levels, decreased SD | normal SD | normal SD | normal SR |
| *pyr1-1pyl1-1 pyl2-1pyl4-1pyl5 pyl8-1* | *pyrpyl112458* | Q169stop, SALK_054640, GT_2864, SAIL_517_C08, SM3_3493, SAIL_1269_A02 | (Gonzalez-Guzman *et al.*, 2012) | ABA insensitivity, increased SD | increased SD | increased SD | increased SR |
| *er-5* | *er-5* | GK-182D08 | isolated in this study | increased SD | increased SD | increased SD | normal SR |
| *er-6* | *er-6* | GK-364C05 | isolated in this study | increased SD | normal SD | increased SD | normal SR |
| *erl1* | *erl1* | GK-109G04 | isolated in this study | normal SD | normal SD | normal SD | normal SR |
| *erl2-2* | *erl2-2* | SALK_015275C | (Nanda *et al.*, 2019) | normal SD | normal SD | normal SD | normal SR |
| *erl2-3* | *erl2-3* | GK-486E03 | (Yang *et al.*, 2022b) | normal SD | normal SD | normal SD | normal SR |
| *sdd1* | *sdd1* | GK-627D04 | (Hara *et al.*, 2007) | increased SD | increased SD | increased SD | normal SR |
| *sdd1-4* | *sdd1-4* | GK-693D08 | isolated in this study | increased SD | increased SD | increased SD | decreased SR |
| *tmm stkr1* | *tmm* | SALK_115723C | isolated in this study, also homozygous for T-DNA insert in *STKR1* (Nietzsche *et al.*, 2018) | increased SD | normal SD | increased SD | decreased SR |

### Experiment 1 - differences between stomatal anatomical traits in different leaves (Figure 2)

Plants were grown in growth cabinets (Percival AR-22L, Percival Scientific, IA, USA) at 10 h light : 14 h dark photoperiod (light intensity 250 μmol m^-2^ s^-1^), relative air humidity 60% during day and 80% during night and temperature 23°C and 19°C during day and night, respectively. Leaves to be sampled for anatomical analyses were numbered according to the protocol of Farmer *et al*. (2013). Leaves 6 and 8 were sampled from plants at 7 weeks of age, and leaves 10 and 12 were sampled as plants had started bolting at 8 weeks of age from different plants grown in parallel. For analyses of stomatal patterning, the sampled leaves were cut in half at the midvein and leaf impressions were collected from both the adaxial and abaxial leaf side, one from each leaf half (Figure 2A) with dental silicone (Speedex light body, Coltene/Whaledent AG; Alstätte, Switzerland and oranwash L, Zhermack; www.zhermack.com for adaxial and abaxial leaf side, respectively) as in Casson *et al*. (2009). Secondary nail varnish imprints were taken from the silicone and transferred to microscope slides with transparent tape. One area of 0.26 mm^2^ from the center (with respect to base and tip) and middle (with respect to midvein and leaf edge) of the leaf from each imprint was photographed under a microscope (Kern OBF 133; Kern & Sohn GmbH; www.kern-sohn.com) at 200x magnification. Stomatal density (stomata mm^-2^, including both mature stomata and stomatal precursor cells) and stomatal ratio (ratio of adaxial and abaxial stomatal densities) were determined from the images using ImageJ software (National Institutes of Health, USA; Schneider et al. 2012). Sample size was 28-32 plants for each genotype resulting from two independently grown batches of plants.

**Figure 2.**
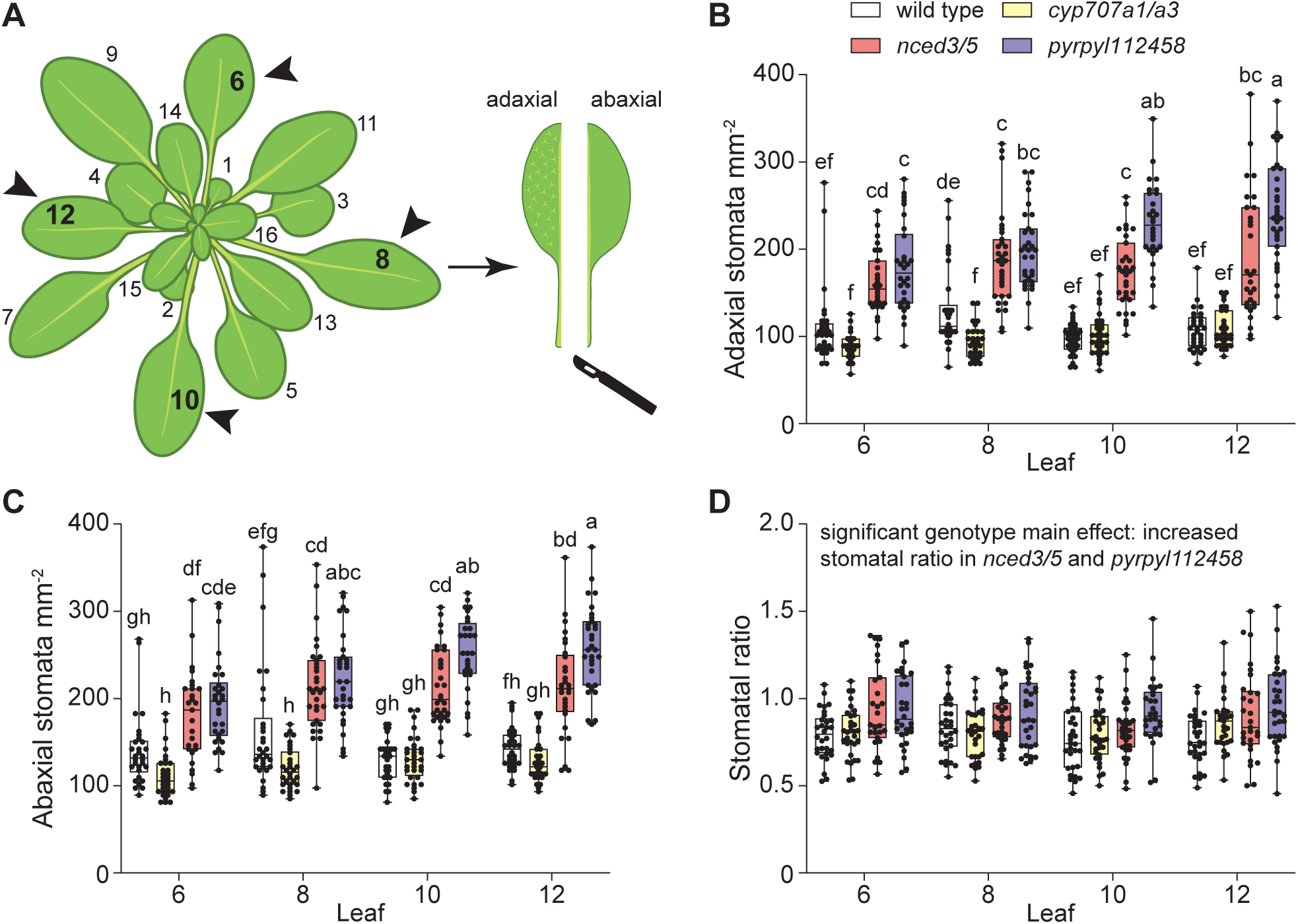
Stomatal density depends on the leaf number in some Arabidopsis genotypes. (A) A scheme depicting the Arabidopsis plant with leaves numbered according to the developmental order. Arrowheads indicate leaves sampled for analyses. Stomatal density in the adaxial (B) and abaxial (C) epidermis, and stomatal ratio (D) in leaves 6, 8, 10, and 12 of selected mutants. The boxes represent the 25th and 75th percentiles, with the median indicated with the horizontal line; the whiskers show the range of values. Solid dots represent individual plants, n = 28-32 plants for each genotype. Significant differences were determined by two-way ANOVA with Tukey post hoc test. Different letters above boxplots represent significant differences at p < 0.05.

### Experiment 2 - stomatal density, index, ratio, and size in leaf 8 of different stomatal development mutants (Figures 3-6)

Plants were grown in a growth cabinet (Microclima Arabidopsis MCA1600-3LP6-E, Snijders Scientific, Tilburg, Netherlands) under the same conditions as in experiment 1, with the exception of uniform relative air humidity of 70% during both day and night. Plants were photographed at 4 weeks of age and the projected rosette area was determined with ImageJ software. Leaves were numbered as in experiment 1 and leaf 8 was sampled from 6-week-old plants. Epidermal imprints were prepared and analyzed as in experiment 1 with some additions. In the first replicate of experiment 2, stomatal density was analyzed from three areas of 0.26 mm^2^ (all in the middle of the leaf with respect to midvein and leaf edge, one area in the center as in experiment 1 and one area each sampled more towards leaf base or tip) and the mean of respective areas was used as the value for respective samples in statistical analyses. As there was no difference in stomatal density between different areas sampled from the same imprint (Supplementary Figure 1), only a single area from leaf center and middle per imprint was analyzed in other experiments and replicates. Stomatal index was calculated as the proportion of stomata from all epidermal cells (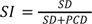, where *SI* is stomatal index, *SD* stomatal density and *PCD* pavement cell density). Size of mature stomata (excluding stomatal precursors) was measured from epidermal imprints using ImageJ software, with length of five stomata per imprint measured and averaged to gain a representative number per sample for subsequent analysis. Sample size was 9-10 plants for each genotype resulting from two independently grown batches of plants.

**Figure 3.**
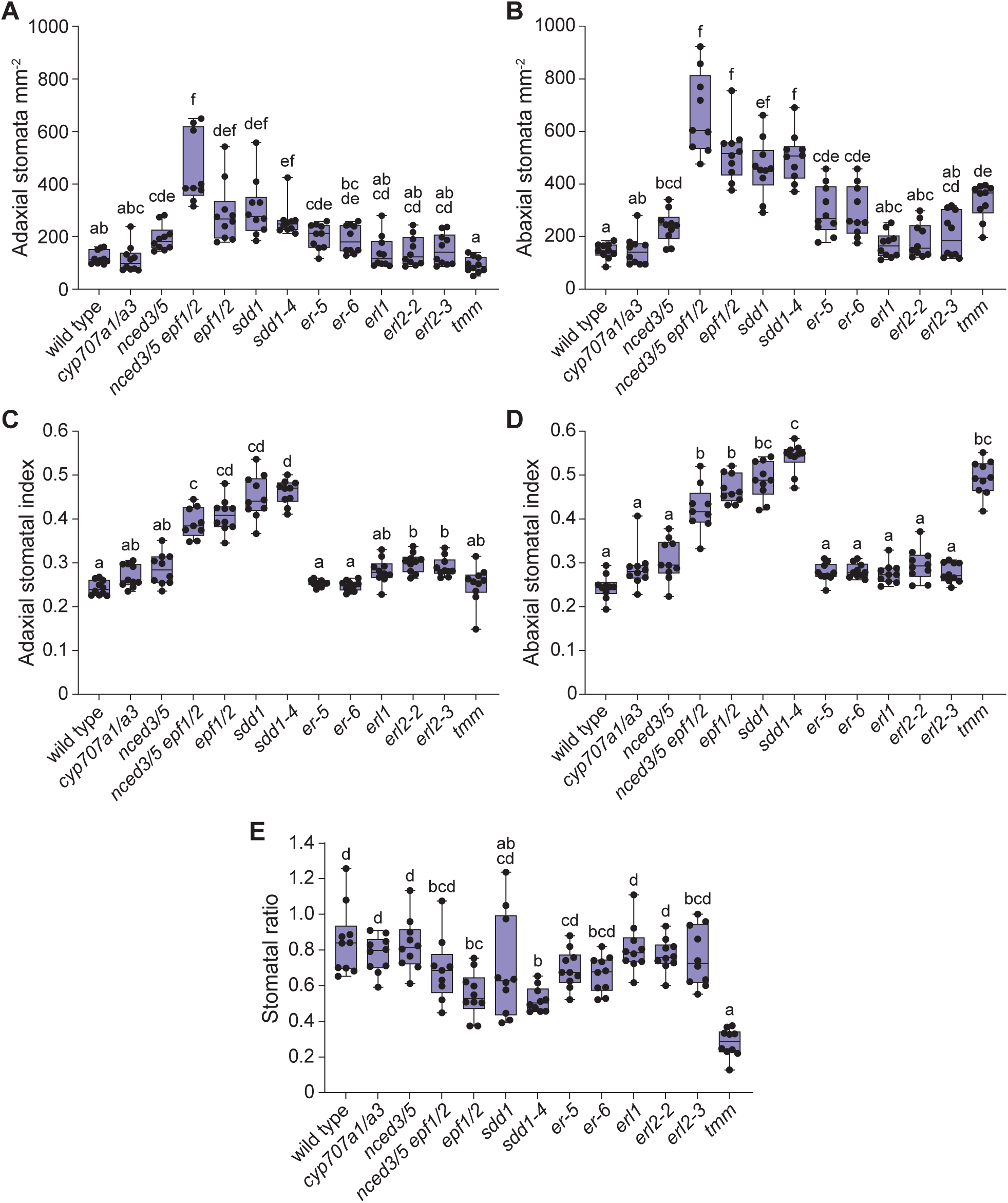
Lack of major regulators of stomatal development differently affects adaxial and abaxial stomatal patterns. Stomatal density in the adaxial (A) and abaxial (B) epidermis, stomatal index in the adaxial (C) and abaxial (D) epidermis, and stomatal ratio (E) of stomatal developmental and ABA biosynthesis or catabolism mutants. The boxes represent the 25th and 75th percentiles, with the median indicated with the horizontal line; the whiskers show the range of values. Solid dots represent individual plants, n = 9-10 plants for each genotype. Significant differences were determined by one-way Welch ANOVA with Dunnett T3 post hoc test. Different letters above boxplots represent significant differences at p < 0.05.

### Statistical analyses

Both experiments were conducted twice with independently grown batches of plants. Pooled data from both batches is shown for both experiments. Statistical analyses were carried out with GraphPad Prism version 10 (GraphPad Software, Boston, MA, USA) or R (version 4.2.1). Two-way ANOVA with Tukey post hoc test, one-way Welch ANOVA with Dunnett T3 post hoc test, or linear regression were used as indicated in figure legends. All effects were considered significant at p < 0.05.

## Results

### Stomatal density depends on leaf number and ABA preferentially suppresses adaxial stomatal development

Most studies addressing stomatal density in true leaves (leaves other than cotyledons) sample an unspecified fully expanded leaf for analysis. As Arabidopsis fully expanded leaves vary in shape and size, it is possible that developmentally distinct leaves also differ in their stomatal patterning. To address the potential within-plant variation in stomatal densities and in the distribution of stomata between adaxial and abaxial leaf surfaces, we analyzed stomatal density and ratio in leaves 6, 8, 10, and 12 in Arabidopsis (Figure 2A). To test if ABA regulates stomatal distribution between adaxial and abaxial leaf surfaces, we included mutants with higher (*cyp707a1/a3*) or lower (*nced3/5*) ABA levels, or impaired ABA perception *(pyrpyl112458*) in our analysis (Table 1).

Both adaxial and abaxial stomatal density was increased in all studied leaves of the *nced3/5* and *pyrpyl112458* mutants compared with wild type plants (Figure 2B, C). A decrease in stomatal density in the *cyp707a1/a3* mutant was significant only in leaf 8 (Figure 2B, C). While stomatal densities of most mutants were similar in different leaves, the ABA perception mutant *pyrpyl112458* had higher adaxial and abaxial stomatal densities in leaves 10 and 12 compared with leaf 6 (Figure 2B, C). Thus, stomatal density depends on leaf number in some Arabidopsis genotypes.

Stomatal ratios were similar across different leaves in all studied mutants (Figure 2D). The ABA biosynthesis mutant *nced3/5* and ABA perception mutant *pyrpyl112458* had slightly higher stomatal ratios compared with Col-0 and *cyp707a1/a3* (significant main effect of genotype, Table 2).

**Table 2.** ANOVA results for Figure 2D.

|  | df | SS | MS | F-value | P-value |
| --- | --- | --- | --- | --- | --- |
| **genotype** | 3 | 1.54 | 0.514 | 15.53 | **< 0.0001** |
| leaf | 3 | 0.15 | 0.051 | 1.55 | 0.20 |
| genotype x leaf | 9 | 0.28 | 0.031 | 0.96 | 0.48 |
| residuals | 465 | 15.39 | 0.033 |  |  |
| total | 480 | 17.37 |  |  |  |

### SDD1, EPF1/2, ERL2, and TMM differently affect stomatal development in adaxial and abaxial epidermis

The increase in stomatal ratio in ABA biosynthesis- or perception-deficient mutants indicates that adaxial and abaxial stomatal development are at least partly independently regulated. We aimed to test whether major regulators of abaxial stomatal development similarly affect adaxial stomatal development. To this end, we obtained T-DNA insertion lines for key components of the stomatal development signaling pathway (Table 1, components shown in bold in Figure 1, representative images of 4 week-old plants shown in Supplementary Figure 2) and analyzed stomatal density, index and size in both adaxial and abaxial epidermis of leaf 8 in these mutants. The components studied in our experiment included the SDD1 protease, the signaling peptides EPF1 and EPF2, and the peptide receptor complex components ER, ERL1, ERL2, and TMM (Figure 1). We also included mutants with lower or higher ABA levels (*nced3/5*, *cyp707a1/a3*) and a mutant combining low ABA levels with lack of EPF1 and EPF2 (*nced3/5 epf1/2*).

Most studied mutations affected stomatal density in adaxial and abaxial epidermis in a similar manner: stomatal density was increased compared with wild type by the *nced3/5*, *epf1/2*, *nced3/5 epf1/2*, *sdd1* and *er* mutations, and not significantly affected by the *cyp707a1/a3*, *erl1* and *erl2* mutations (Figure 3A, 3B, Table 1). Stomatal density was increased only in the abaxial epidermis in the *tmm* mutant (Figure 3A, 3B, Table 1). Stomatal index was higher than in wild type in both adaxial and abaxial epidermis in the *epf1/2*, *nced3/5 epf1/2*, *sdd1*, and *sdd1-4* mutants (Figure 3C, 3D), whereas it was higher only in the adaxial epidermis in both studied *erl2* mutants (Figure 3C), and higher only in the abaxial epidermis in the *tmm* mutant (Figure 3D). Stomatal index was not significantly affected in the *cyp707a1/a3* and *nced3/5* mutants (Figure 3C, 3D). Stomatal ratio was significantly lower than in wild type in the *epf1/2*, *sdd1-4* and *tmm* mutants (Figure 3E).

Stomatal size varied very little between different plant lines: stomatal length of studied mutants was similar to wild-type in both adaxial and abaxial leaf surface, with the exception of *sdd1-4* that had smaller stomata than wild-type plants in abaxial epidermis (Figure 4A, 4B). We found no significant relationship between stomatal density and length on the adaxial leaf surface (Figure 4C), whereas there was a significant negative linear relationship between stomatal density and length in abaxial epidermis only when the *nced3/5 epf1/2* mutant was excluded from analysis (Figure 4D).

**Figure 4.**
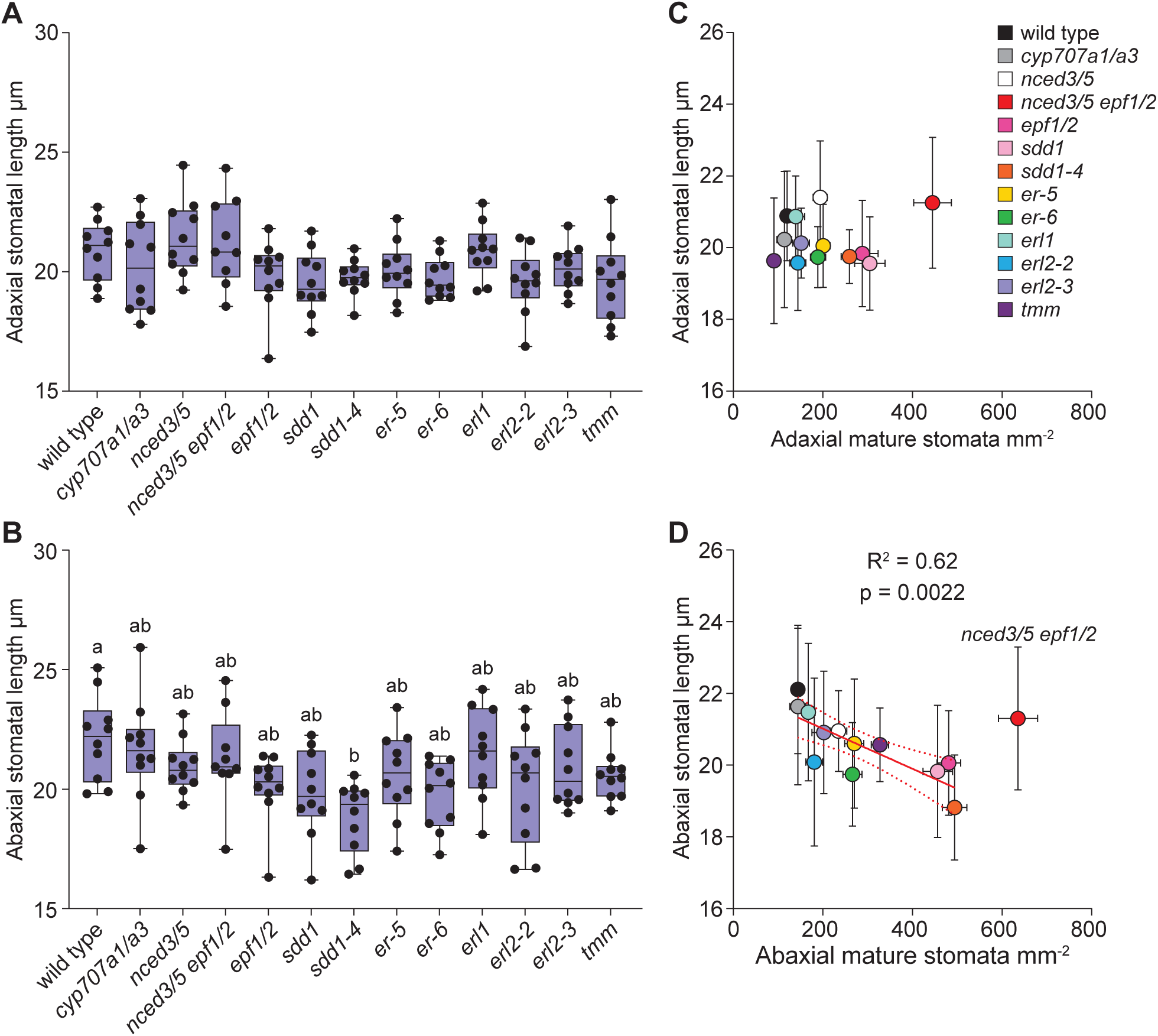
The size of stomata varies very little in adaxial and abaxial leaf sides. Length of stomata in the adaxial (A) and abaxial (B) leaf side. Relationship between stomatal density and stomatal length in adaxial (C) and abaxial (D) leaf side. The boxes represent the 25th and 75th percentiles, with the median indicated with the horizontal line; the whiskers show the range of values. Solid dots represent individual plants, n = 9-10 plants for each genotype. Significant differences were determined by one-way Welch ANOVA with Dunnett T3 post hoc test (A, B), linear regression was used in (C, D). Different letters above boxplots represent significant differences at p < 0.05. The slope of the linear regression differs significantly from zero (p<0.05) only in the abaxial side when the *nced3/5 epf1/2* quadruple mutant is excluded (D).

### Stomatal development rarely arrests in the precursor state in adaxial epidermis

In plant lines with aberrant stomatal development, stomata sometimes arrest in a precursor state. In most studied mutants, we did not detect more stomatal precursors than in wild type plants (Supplementary Figure 3). When stomatal precursors were found in fully expanded leaf 8, such cells usually only occurred in the abaxial epidermis. This was true for the *epf1/2*, and *er* mutants, whereas we found stomatal precursors in both adaxial and abaxial epidermis in the *nced3/5 epf1/2* mutant (Figure 5). Thus, stomatal development rarely arrests in the precursor state in the adaxial epidermis.

**Figure 5.**
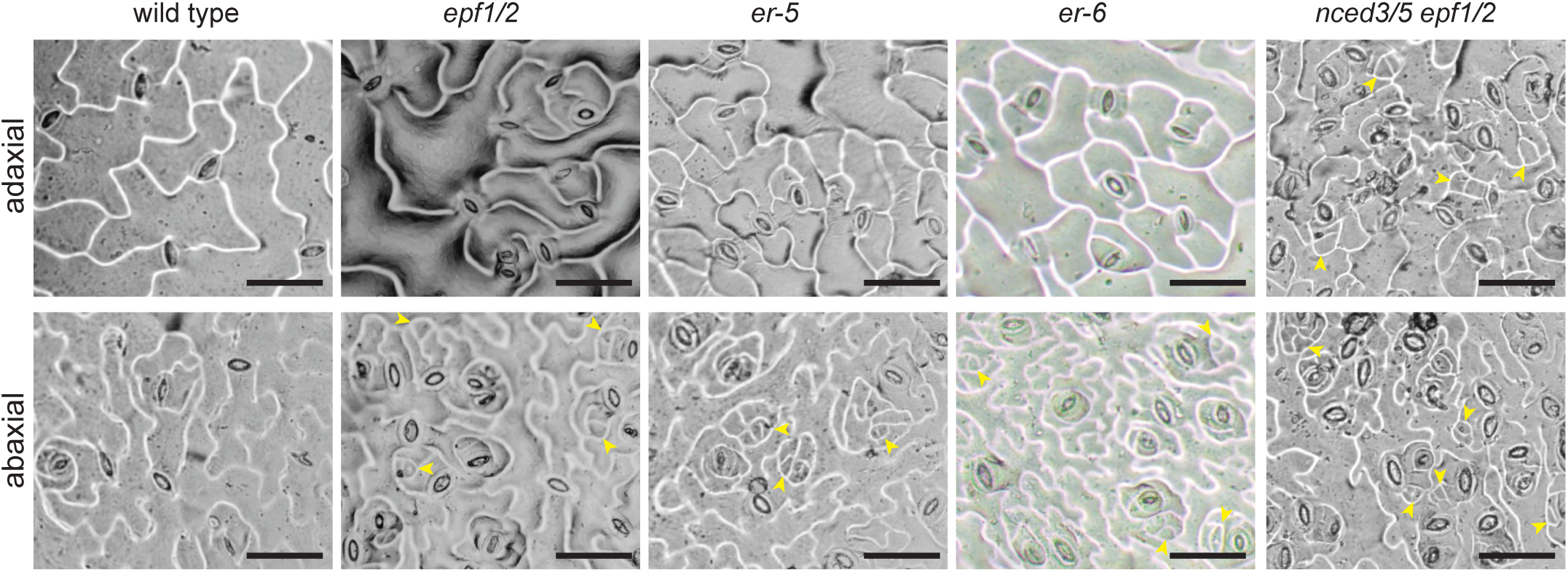
Stomatal precursors are mostly found in the abaxial epidermis. Representative images of adaxial and abaxial stomatal impressions of analyzed mutants that harbor stomatal precursors in fully expanded leaves. Yellow arrowheads indicate stomatal precursors. The scale bar represents 50 µm.

### Stomatal density is negatively related with plant growth

Total stomatal density (sum of adaxial and abaxial stomatal densities) was significantly increased in the *nced3/5*, *nced3/5 epf1/2*, *epf1/2*, *sdd1*, *er* and *tmm* mutants, with the most prominent effects in *epf1/2*, *sdd1* and *epf1/2 nced3/5* mutants (Figure 6A). The projected rosette area was significantly reduced in the mutant with the highest stomatal density – the *nced3/5 epf1/2* double mutant (Figure 6B). Across all studied mutants, there was a significant negative relationship between stomatal density and plant size (Figure 6C).

**Figure 6.**
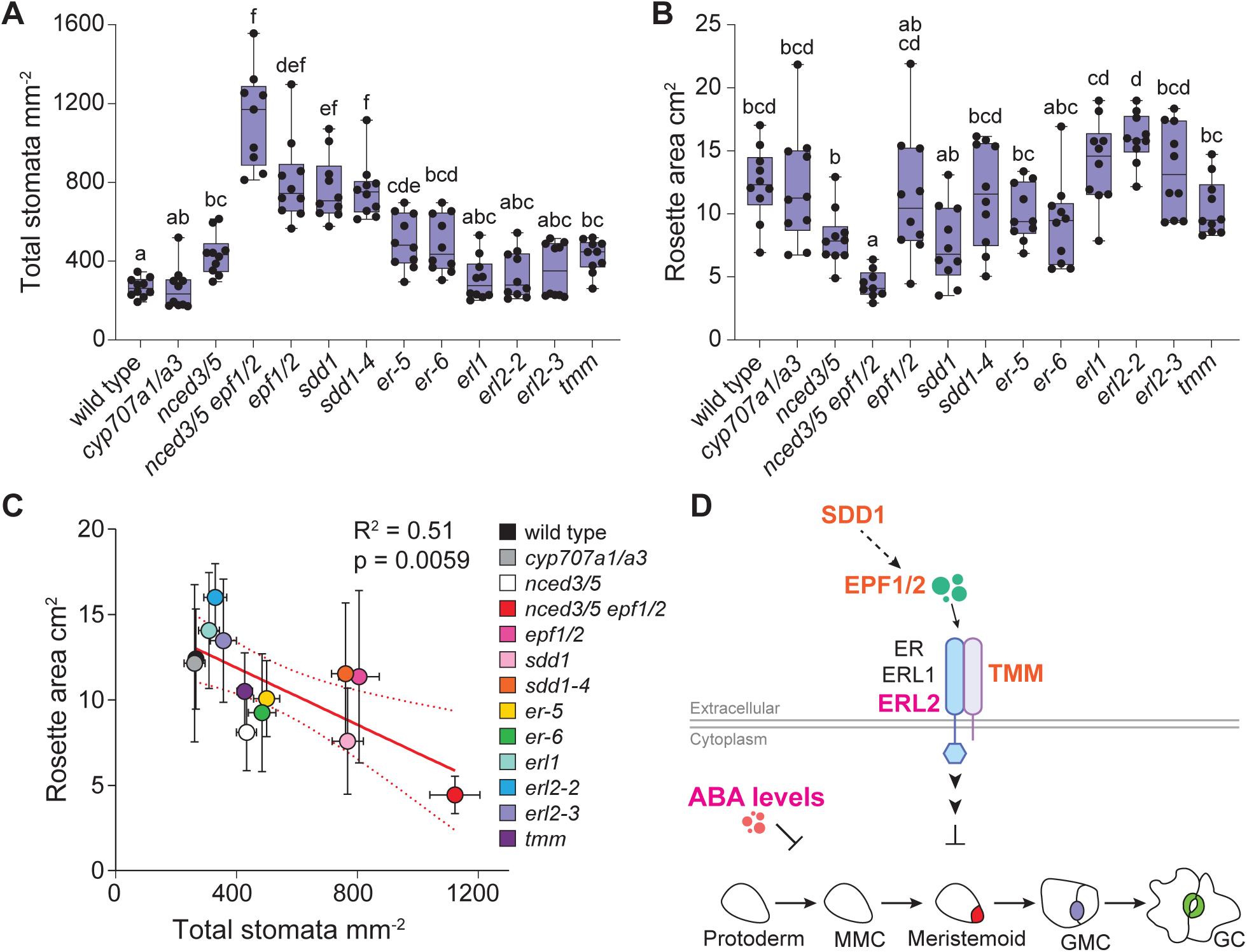
Stomatal density is negatively related with plant growth. Total stomatal density (A) and projected rosette area (B) of stomatal developmental and ABA biosynthesis or catabolism mutants. (C) Relationship between total stomatal density and rosette area. (D) A model based on the obtained results suggests that SDD1, EPF1/2, and TMM are more important for suppressing stomatal development in the abaxial epidermis (indicated by orange color), whereas ERL2 and ABA suppress stomatal development preferentially in the adaxial epidermis (indicated in pink). The boxes represent the 25th and 75th percentiles, with the median indicated with the horizontal line; the whiskers show the range of values. Solid dots represent individual plants, n = 9-10 for each genotype. Significant differences were determined by one-way Welch ANOVA with Dunnett T3 post hoc test (A, B) and linear regression was used in (C). Different letters above boxplots represent significant differences at p < 0.05.

## Discussion

Here we addressed within-plant variation in stomatal density and ratio in Arabidopsis and the role of major stomatal developmental regulators in stomatal patterning in adaxial and abaxial epidermis. We show that while stomatal ratio is similar across developmentally different leaves, stomatal density in some Arabidopsis genotypes depends on the sampled leaf (Figure 2). We also found that a lack of certain stomatal development regulators leads to different developmental outcomes in upper and lower leaf surfaces (Figures 3-5), indicating their different importance in adaxial and abaxial stomatal development. Signaling components that preferentially affect stomatal development in adaxial or abaxial leaf surface are indicated in pink or orange, respectively, in the model in Figure 6D.

In wild type Arabidopsis, stomatal density was similar in developmentally different fully expanded leaves, whereas in the ABA perception mutant *pyrpyl112458*, stomatal densities differed between distinct fully expanded leaves (Figure 2B, C). Thus, to reduce variation in stomatal density analyses in true leaves, we advise sampling a defined fully expanded leaf for analyses of stomatal patterning. Previous studies have shown heterogeneity in stomatal density as well as stomatal conductance across the leaf (Tichá, 1982; Weyers and Lawson, 1997; Sweet *et al*., 2017). Our results suggest that when sampling stomatal density from the middle (with respect to midvein and leaf edge) areas of the Arabidopsis leaf, there is very little heterogeneity in stomatal density (Supplementary Figure 1), making Arabidopsis a good model for studies on stomatal density in both adaxial and abaxial epidermis.

Stomatal ratio did not differ between leaves in any of our studied genotypes (Figure 2D). However, the stomatal ratio values that we measured in true leaves of wild-type plants were higher than reported before in cotyledons (Geisler *et al*., 1998; Berger and Altmann, 2000; Delgado *et al*., 2019), indicating that there is within-plant variation also in leaf stomatal ratio. Differences between stomatal characteristics depending on leaf number have been discussed before, stressing more severe effects of different mutations on stomatal development in cotyledons than in true leaves (Dow *et al*., 2014). Understanding how stomatal development between cotyledons and true leaves differs is important for practical applications aiming to regulate plant water use efficiency or productivity via modulation of stomatal numbers, as cotyledons have a minor role in overall plant gas-exchange and physiology.

The lack of negative regulators of stomatal development increased stomatal density more strongly in the abaxial epidermis, leading to reduced stomatal ratios in *epf1/2*, *sdd1-4* and *tmm* mutants (Figure 3). Similarly reduced stomatal ratios have been found before for these mutants (Berger and Altmann, 2000; Dow *et al*., 2014; Franks *et al*., 2015; Hronková *et al*., 2015; Vráblová *et al*., 2017), indicating a more prominent role of respective signaling components in adjusting abaxial than adaxial stomatal development. More severely disrupted abaxial than adaxial stomatal patterns were also found in plants with modified expression levels of SPCH (Dow *et al*., 2014), a repressor of SPCH (IDD16, Qi *et a*l., 2019), or STOMAGEN/EPFL9 (Hronková *et al*., 2015). As studies of stomatal form and function have largely focused on abaxial stomata, it is expected that the currently identified stomatal developmental signaling components are of major importance in abaxial epidermal patterning. Focusing on the adaxial stomatal development in the future might reveal new signaling components or pathways in stomatal development. Several studies indicate that adaxial stomatal development is more sensitive to changes in environmental conditions, such as light levels (Hronková *et al*., 2015) or relative air humidity (Devi and Reddy, 2018; Tulva *et al*., 2023, Preprint). Between-species comparisons also suggest that adaxial stomatal numbers have been adjusted more freely during evolution (Muir *et al*., 2023). The presence of adaxial stomata in amphistomatic leaves has been associated with improved CO_2_ uptake (Parkhurst and Mott, 1990; Drake *et al*., 2019), photosynthesis (Xiong and Flexas, 2020) and faster stomatal responsiveness (Haworth *et al*., 2018). Understanding how adaxial stomatal development is adjusted in plants is necessary to better understand the physiology of amphistomatous plants.

We found that the lack of ERL2 alone was sufficient to increase adaxial but not abaxial stomatal index (Figure 3), indicating that ERL2 is more involved in suppressing adaxial than abaxial stomatal development. Indeed, similarly increased adaxial but not abaxial stomatal index was found before in the *erl1erl2* double mutant (Kong *et al*., 2012). Our findings are in line with previous evidence showing distinct functions of specific ER family proteins in fine-tuning stomatal development. The specific roles of ER, ERL1 and ERL2 manifest in their different expression patterns (Shpak *et al*., 2004), binding of different EPFs (Lee *et al*., 2012), and different regulation of their subcellular localization (Yang *et al*., 2022*b*). Our findings indicate that specific ER family protein members also contribute to a different extent to stomatal differentiation in adaxial and abaxial epidermis.

Stomatal precursor cell formation was shown to cease earlier in adaxial compared with abaxial epidermis in cotyledons (Geisler and Sack, 2002). Here, stomatal precursor cells, if present in fully expanded leaves of stomatal development mutants, were typically found only in the abaxial epidermis (Figure 5). This is in line with prolonged stomatal initiation in abaxial epidermis. As an exception, the *nced3/5 epf1/2* mutant had arrested stomatal precursor cells both in the adaxial and abaxial epidermis of fully expanded leaves (Figure 5). Thus, low ABA levels in the *epf1/2* mutant background led to increased numbers of stomatal precursors in the adaxial epidermis, indicating that normal ABA levels can rescue stomatal differentiation in the adaxial epidermis of the *epf1/2* mutant that only harbors stomatal precursors in the abaxial epidermis. The asymmetry of the presence of cells arrested in a stomatal precursor state between adaxial and abaxial epidermis indicates that different mechanisms control stomatal differentiation in adaxial and abaxial epidermis.

ABA is considered a suppressing signal for stomatal development largely based on studies of ABA biosynthesis and catabolism mutants. We found higher stomatal densities in *nced3/5* and *pyrpyl112458* (Figure 2), in line with generally increased abaxial stomatal density of ABA biosynthesis and perception mutants (Chater *et al*., 2015; Hsu *et al*., 2018; Jalakas *et al*., 2018; Merilo *et al*., 2018). However, we found no effect of the *cyp707a1/a3* mutations on stomatal density in most leaves (Figure 2-3) and no effect on stomatal index (Figure 3). This difference between our and previous studies (Tanaka *et al*., 2013*a*; Jalakas *et al*., 2018) might in part be explained by variable ABA levels in ABA catabolism mutants under different growth conditions as has been found before for some ABA biosynthesis mutants (Merilo *et al*., 2018). Although the ABA-deficient *nced3/5* mutant had higher stomatal density in our experiments, there was no change in stomatal index, so the change in density might result from the effects of ABA-deficiency on cell growth rather than stomatal development. Stomatal ratio was slightly increased in plants deficient in ABA biosynthesis or perception (Figure 2D, Table 2), suggesting that ABA as a negative regulator of stomatal development might preferentially suppress stomatal development in the adaxial epidermis. Together, our data suggest that in true leaves, ABA levels have a minor effect on the proportion of cells entering the stomatal lineage and on the distribution of stomata between leaf surfaces, whereas mutations in stomatal development pathway components have much larger effects. Studies finding a role for ABA levels in regulating stomatal index have often been carried out in cotyledons of plate-grown seedlings subjected to 100% relative air humidity (Tanaka *et al*., 2013*a*; Mohamed *et al*., 2023). Leaf ABA levels depend on air humidity (McAdam *et al*., 2016) and low relative air humidity primes ABA sensitivity in leaves (Pantin *et al*., 2013). Thus, it is expected that the quantitative effects of ABA on stomatal patterning are different between plants that have not been subjected to naturally fluctuating air humidity and those that have. Our experiments suggest that in true leaves grown under moderate relative air humidity, modification of ABA levels in the range achieved by the ABA biosynthesis and catabolism mutants is of minor importance in regulating entry into the stomatal lineage.

A negative relationship between stomatal density and size attributed to packing constraints has been found both across species (Franks and Beerling, 2009) and in Arabidopsis (Dittberner *et al*., 2018). We found no significant relationship between stomatal density and size in the adaxial leaf surface (Figure 4C), whereas there was a significant negative relationship between stomatal density and size in abaxial leaf surface when excluding the extreme mutant *nced3/5 epf1/2* (Figure 4D). Conversely, we recently found a weak negative relationship between stomatal density and size in adaxial, but not abaxial epidermis in a different set of Arabidopsis mutants (Tulva *et al*., 2023, Preprint). Together, our data suggest that stomatal size and density are rather weakly linked in Arabidopsis. The large stomatal size combined with high stomatal density in the *nced3/5 epf1/2* (Figure 4D) indicates that it is possible to break the negative relationship between stomatal size and density and fit many large stomata onto the leaf surface. Thus, spatial constraint to stomatal packing may not be the reason for the often observed negative relationship between stomatal density and size.

We previously found a negative relationship between plant growth and stomatal density in plant lines, where stomatal density was altered by the *epf1* and/or *epf2* mutations (Tulva *et al*., 2023, Preprint). Here, we show that this relationship is more universal, as it holds for a set of plant lines with varying stomatal density due to different mutations (Figure 6C). Despite the improved CO_2_ uptake capacity and increased photosynthesis found before in high stomatal density mutants (Schlüter *et al*., 2003; Tanaka *et al*., 2013*b*; Tulva *et al*., 2023, Preprint), such mutants fail to achieve better growth. Possibly, the resources obtained by increased net assimilation are used to carry out expensive cell divisions (Dow *et al*., 2014; Franks *et al*., 2015; Bowers, 2018), or for investment into roots to improve water uptake capacity, as discussed before (Hepworth *et al*., 2016). Future studies of above- and below-ground growth in plants with altered stomatal conductance due to different stomatal apertures or densities are needed to test the latter hypothesis.

In conclusion, stomatal patterning in Arabidopsis is at least partly differently controlled in the adaxial and abaxial epidermis. In wheat, adaxial stomata are responsible for the majority of leaf gas-exchange, they are more responsive to light than abaxial stomata, and adaxial stomatal density is higher and more responsive to growth at elevated CO_2_ levels (Wall *et al*., 2022, 2023). These recent works show that adaxial stomata are critical for gas-exchange in cereals and, given the relatively large stomatal ratios in Arabidopsis (Figure 3), can potentially account for a large proportion of gas-exchange also in herbs. Stomatal density, size and shape impact on photosynthesis, the same is likely true for stomatal distribution between upper and lower surfaces in leaves (Drake *et al*., 2019; Harrison *et al*., 2020). Cuticular conductance is positively related with stomatal density, whereas adaxial and abaxial leaf surfaces can have different cuticular conductances (Márquez *et al*., 2022). Understanding how adaxial stomata and stomatal ratio are determined and how they affect plant gas-exchange, including net assimilation, stomatal and cuticular conductances, is needed to develop crops that optimally balance water use with CO_2_ assimilation.

## Acknowledgements

We thank Agnes Vaht, Marilin Poogen and Tamta Jashi for help with plant growth and sample collection, Elfi Aliyeva for help with sample imaging, and Kārlis Dieviņš for help with projected rosette area measurements.

## Author contributions

Conceptualization - P.J., I.T., H.H., Data curation and analysis - all authors; Funding Acquisition - H.H.; Investigation - all authors; Supervision - I.T., H.H.; Visualization - P.J.; Writing – Original Draft - P.J., H.H.; Writing – Review & Editing - all authors

## Data availability

Primary data for figures and tables presented in this manuscript are included as supplementary material.

