## Supplementary Figures for "Stomatal patterning is differently regulated in adaxial and abaxial epidermis in Arabidopsis"

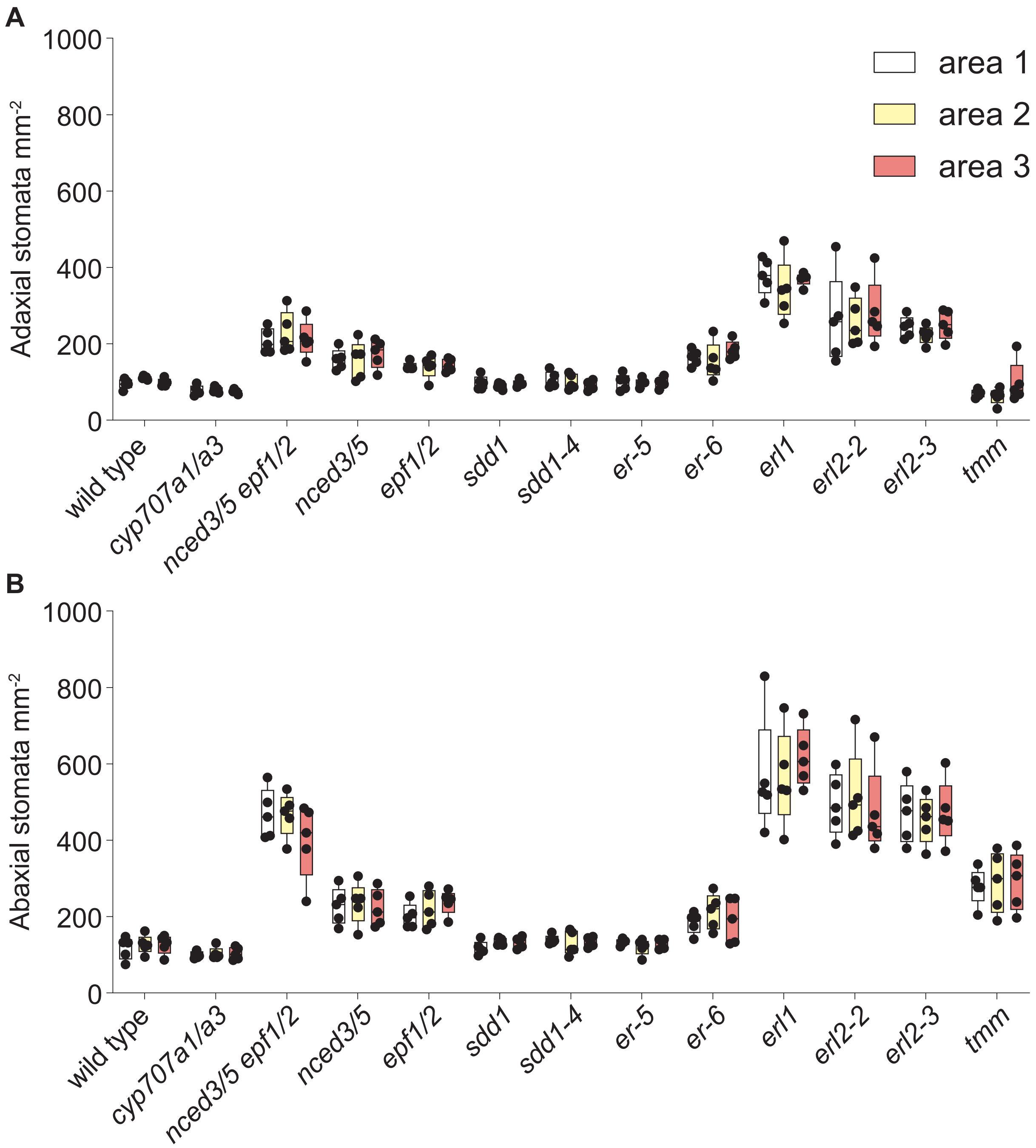


**Supplementary Figure 1.** Comparison of stomatal density in three different areas sampled from the same imprint in the adaxial (A) and abaxial (B) leaf side. The boxes represent the 25th and 75th percentiles, with the median indicated with the horizontal line; the whiskers show the range of values. Solid dots represent individual plants, n = 5. Two-way ANOVA with genotype and area as factors showed no significant interaction or effect of sampled area on stomatal density, only the effect of genotype was significant.

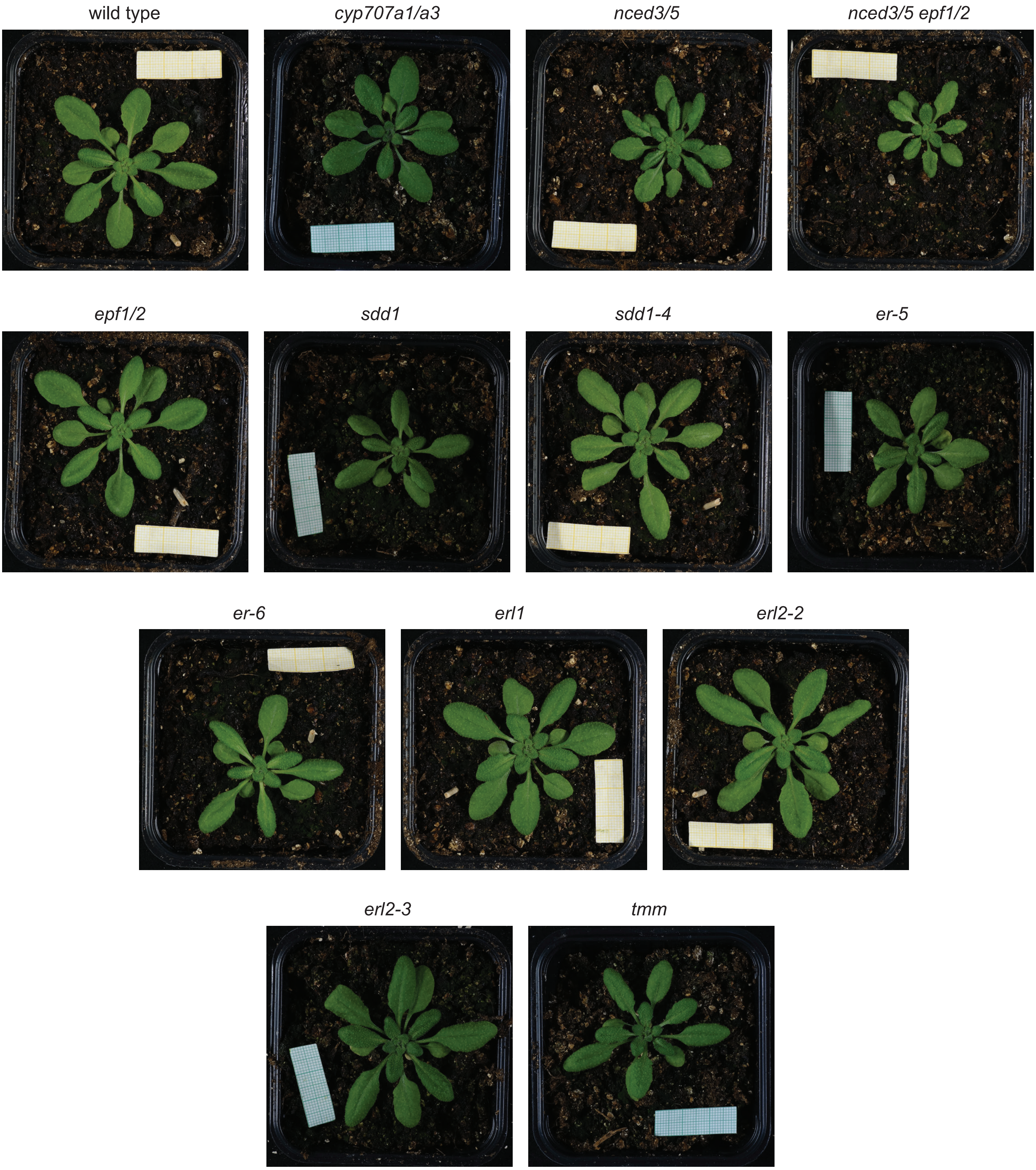


**Supplementary Figure 2.** Representative images of 4-week-old plants.

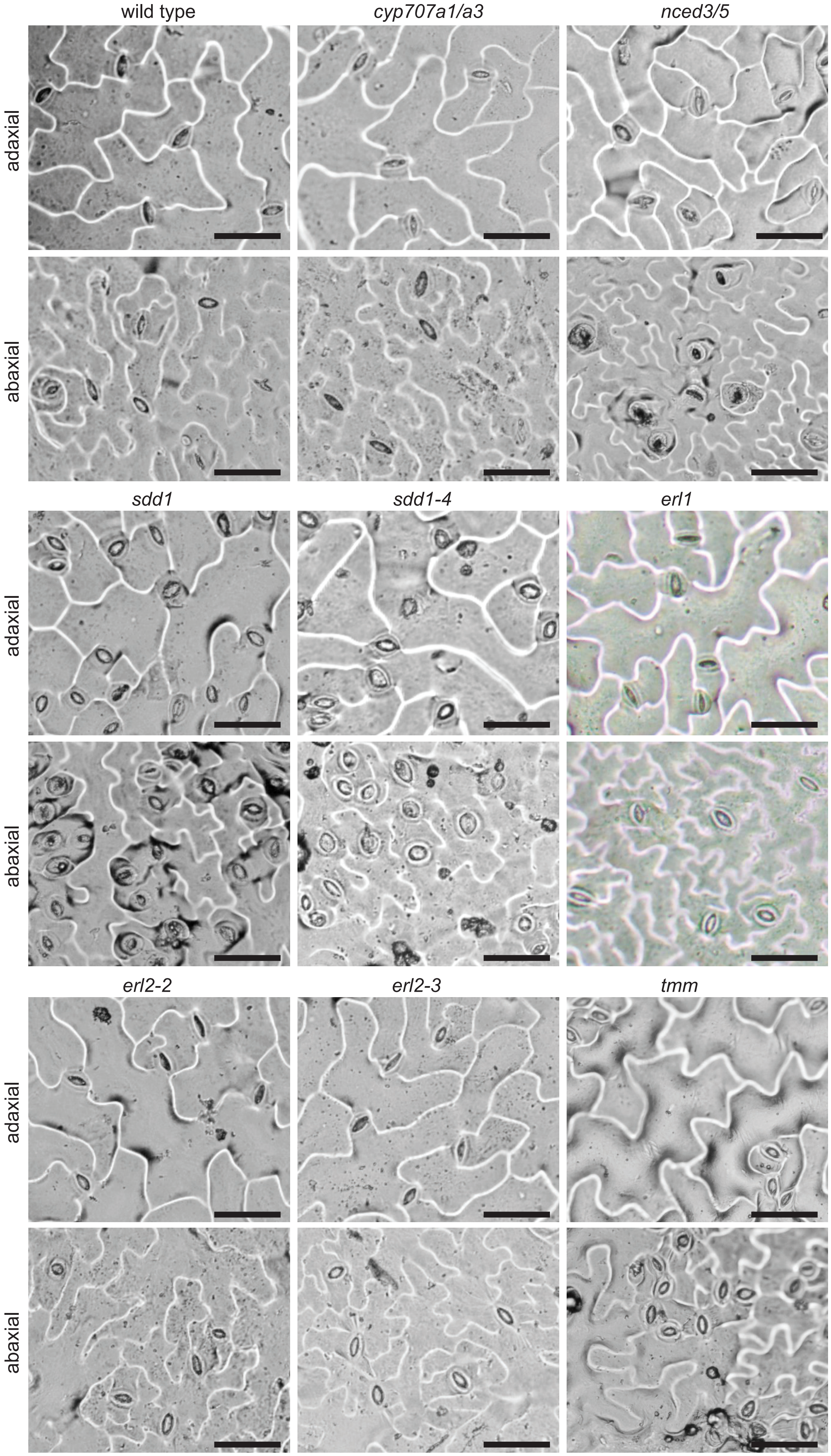


**Supplementary Figure 3.** Representative images of abaxial and adaxial stomatal impressions. The scale bar represents 50 µm.
