## Supplementary Table 1 for "Stomatal patterning is differently regulated in adaxial and abaxial epidermis in Arabidopsis"

**Supplementary Table 1.** Primers used for genotyping studied mutants

| **Allele name** | **Line name** | **Border primer** | **Left primer (LP)** | **Right primer (RP)** | **Genotyping principle** |
| --- | --- | --- | --- | --- | --- |
| *nced3-2* | GK-129B08 | ATATTGACCATCATACTCATTGC | GTCAGCCACGAGAAGCTACAC | TTCACCGGTTTTGAGATTCAG | Product with LP + RP from wild-type allele, and with border primer + RP from mutant allele |
| *nced5-2* | GK-328D05 | ATATTGACCATCATACTCATTGC | TAACACCAAACCCAACCAAAC | TGACTCAACCCAAACCATCTC | Product with LP + RP from wild-type allele, and with border primer + RP from mutant allele |
| *epf1-1* | SALK_137549 | ATTTTGCCGATTTCGGAAC | GGTGCATGTTCGACACTCTTC | CATGGTCATGTCCCGGAGAAGC | Product with LP + RP from wild-type allele, and with border primer + RP from mutant allele |
| *epf2-2* | GK-673E01 | ATATTGACCATCATACTCATTGC | ATGACGAAGTTTGTACGCAAG | AATCTGATTCGGTCGTGTC | Product with LP + RP from wild-type allele, and with border primer + RP from mutant allele |
| *cyp707a1-1* | SALK_069127 | ATTTTGCCGATTTCGGAAC | CATGAACGTATTGGGTTTTGG | TCCTGATATTGAATCCATCGC | Product with LP + RP from wild-type allele, and with border primer + RP from mutant allele |
| *cyp707a3* | SALK_101566 | ATTTTGCCGATTTCGGAAC | GTTCCTGGAAGATTAATCGGC | ACGTGCTCTCGTCACTCTCTC | Product with LP + RP from wild-type allele, and with border primer + RP from mutant allele |
| *pyr1-1* | Q169stop |  | TCGGTTCGAGAAAGAGAATCG | CGTCATTCTCATCATAAGAAAATGG | PCR product digested with HpyCH4V (New England Biolabs), cuts wild-type but not mutant product |
| *pyl1-1* | SALK_054640 | ATTTTGCCGATTTCGGAAC | TGCCAATTTTCAGACATTAAGC | AACCATGCCTTCCGATTTAAC | Product with LP + RP from wild-type allele, and with border primer + RP from mutant allele |
| *pyl2-1* | GT_2864 | CCGTTTACCGTTTTGTATATCCCG | ATGAGCTCATCCCCGGCCG | TTCATCATCATGCATAGGTGCAG | Product with LP + RP from wild-type allele, and with border primer + RP from mutant allele |
| *pyl4-1* | SAIL_517_C08 | GCCTTTTCAGAAATGGATAAATAGCCTTGCTTCC | TTCCAATCGTTCCAAATATCG | TAAGACTCGACAACGACGGTC | Product with LP + RP from wild-type allele, and with border primer + RP from mutant allele |
| *pyl5* | SM3_3493 | TACGAATAAGAGCGTCCATTTTAGAGTGA | AAACACAAAGCCTTCACATCC | AAGTTTTGTGAATCCCCCAAC | Product with LP + RP from wild-type allele, and with border primer + RP from mutant allele |
| *pyl8-1* | SAIL_1269_A02 | GCCTTTTCAGAAATGGATAAATAGCCTTGCTTCC | AGAGAGTGGAACCCCATGATC | TTCTTCTTCTTCCTTCATGCG | Product with LP + RP from wild-type allele, and with border primer + RP from mutant allele |
| *er-5* | GK-182D08 | ATATTGACCATCATACTCATTGC | AGCAAAAGATGCACAAAGAGG | CCTGATCATCTGAGCTCTTGC | Product with LP + RP from wild-type allele, and with border primer + RP from mutant allele |
| *er-6* | GK-364C05 | ATATTGACCATCATACTCATTGC | AATATCAAAGGTCCAATCCCG | ATGCACAATACCAAAACCTGC | Product with LP + RP from wild-type allele, and with border primer + RP from mutant allele |
| *erl1* | GK-109G04 | ATATTGACCATCATACTCATTGC | TTTCCAATCATGATGTTGCAG | CAAACAATTGCTCCAGCTTTC | Product with LP + RP from wild-type allele, and with border primer + RP from mutant allele |
| *erl2-2* | SALK_015275C | ATTTTGCCGATTTCGGAAC | AATGACACATCGCTGAGAAGG | TATCTCCATGGCAACAAGCTC | Product with LP + RP from wild-type allele, and with border primer + RP from mutant allele |
| *erl2-3* | GK-486E03 | ATATTGACCATCATACTCATTGC | TATCTCCATGGCAACAAGCTC | AATGACACATCGCTGAGAAGG | Product with LP + RP from wild-type allele, and with border primer + RP from mutant allele |
| *sdd1* | GK-627D04 | ATATTGACCATCATACTCATTGC | TCTTTTGTTGCTGAAAAAGGC | ACACGGTGTCCTCTGATGAAG | Product with LP + RP from wild-type allele, and with border primer + RP from mutant allele |
| *sdd1-4* | GK-693D08 | ATATTGACCATCATACTCATTGC | GTTGAATCTCTTGCGGAAATG | TCTTTTGTTGCTGAAAAAGGC | Product with LP + RP from wild-type allele, and with border primer + RP from mutant allele |
| *tmm* | SALK_115723C | ATTTTGCCGATTTCGGAAC | ATCTAGGGCCCAACACAAGAC | AATTGGTTGAGCCGGTTAATC | Product with LP + RP from wild-type allele, and with border primer + RP from mutant allele |
| *stkr1* | SALK_115723C | ATTTTGCCGATTTCGGAAC | TTGAGAAAAGTATGGCCAAGTG | AAGCTTGGCGGAAAATCTAAG | Product with LP + RP from wild-type allele, and with border primer + RP from mutant allele |
